# Modified OspA Delivered Via Distinct Vehicles and Routes Confers Equivalent Protection Against Tick-Transmitted *Borrelia burgdorferi*

**DOI:** 10.64898/2026.03.03.709130

**Authors:** Suman Kundu, Greg Joyner, Olifan Abil, Maria J. Sanches, Maria Cristina Gingerich, Kieran Blake Holloman, Hong Jin, Biao He, Maria Gomes-Solecki

## Abstract

Vaccines targeting outer surface protein A (OspA) of *Borrelia burgdorferi* protect against Lyme disease by eliciting antibodies that neutralize spirochetes in the *Ixodes scapularis* tick midgut during engorgement before transmission occurs. We evaluated whether different delivery vehicles and administration routes for a modified OspA construct (OspA_BPBPk_) differed in protective efficacy. Four groups of nine C3H-HeN mice were immunized with OspA_BPBPk_ delivered intranasally by live parainfluenza virus 5 (PIV5- OspA_BPBPk_) or by subcutaneous administration of recombinant protein adjuvanted with aluminum hydroxide (rOspA_BPBPk_) using three prime-boost regimens and a vector/adjuvant control: homologous intranasal PIV5-OspA_BPBPk_ (IN/IN), homologous subcutaneous rOspA_BPBPk_ (SC/SC), and heterologous intranasal PIV5-OspA_BPBPk_ prime/subcutaneous rOspA_BPBPk_ boost (IN/SC). Mice were challenged on day 90, approximately 2 months after the boost, with nymphal ticks infected with 19 strains of *B. burgdorferi*. All OspA_BPBPk_-containing vaccines elicited high antigen-specific IgG antibody titers (geometric mean titers of 10⁴–10⁶, approximately 4 log₁₀ higher than the control group), reduced *B. burgdorferi* loads in engorged nymphal ticks by 1.7-2 Log_10_, neutralized *B. burgdorferi* motility in multi-strain cultures, and prevented recovery of viable spirochetes from tissues after tick challenge. While one mouse (1/9, 11%) in the heterologous IN/SC vaccinated group had detectable *flaB* DNA in tissues and seroreactivity to *B. burgdorferi* PepVF, culture of spirochetes from heart and bladder showed no evidence of active infection. Although homologous immunization produced the most consistent results in mice, different delivery vehicles and routes of immunization with OspA_BPBPk_ vaccines did not affect overall efficacy as assessed by recovery of viable *B. burgdorferi* from tissues.

## Introduction

Despite effective antibiotic therapy for most cases of Lyme disease, delayed diagnosis and persistent morbidity underscore the need for vaccines that prevent transmission and/or infection (1, 2). The first generation of Lyme vaccines, based on outer surface protein A (OspA) of *B. burgdorferi*, demonstrated that high-level protection against Lyme disease (76–92%) is achievable in humans (3, 4). Post-licensure surveillance and subsequent analyses confirmed safety (5–7) and several companies continued development of second-generation OspA-based vaccines using recombinant protein (8, 9), mRNA (10) and viral vector (11) technologies.

OspA is an arthropod-phase lipoprotein first identified as a dominant outer membrane lipoprotein expressed by *B. burgdorferi* (12) in the unfed tick midgut (13). Early rodent and *in vitro* studies showed that OspA-specific antibodies can clear *B. burgdorferi* from feeding ticks and inhibit growth in culture in a titer-dependent manner (14–16). Subsequent work in mice and nonhuman primates confirmed that OspA vaccination prevents tick-transmitted infection (13, 17), and that protection was maintained even under immunosuppression (17), consistent with the complete or near complete protective immunity mediated by circulating antibodies. Collectively, these data established OspA as a transmission-blocking immunogen whose efficacy depends predominantly on the magnitude of the mammalian host serologic IgG specific to OspA, at the time of tick feeding. Recent analyses of OspA serology in clinical cohorts and experimental models further support the interpretation that sustained high-titer anti-OspA response is a key correlate of transmission-blocking protection (11, 18, 19).

Recombinant viral vectors such as parainfluenza virus 5 (PIV5) provide an alternative delivery system for vaccines. PIV5 is a non-segmented, negative-strand RNA virus that replicates exclusively in the cell cytoplasm, minimizing the risk of genomic integration, and has an extensive safety record through decades of use in licensed canine vaccines without evidence of zoonotic disease (20, 21). PIV5 tolerates insertion of foreign genes and has been developed as a vector for influenza, RSV, MERS-CoV, SARS-CoV-2, and other pathogen immunogens, where intranasal or parenteral administration elicits strong immune responses after one or two doses, with documented genetic stability and efficacy across species (22–27). Importantly, recent clinical and nonhuman primate studies of intranasal PIV5-vectored SARS-CoV-2 (CVXGA1) and RSV (BLB201) vaccines report favorable safety and immunogenicity profiles (28–30), supporting the translational potential of this vector.

Our interest in developing Lyme disease vaccines that would be acceptable to the public led to the redesign of OspA (OspA_BPBPk_) (11, 31) and development of needle-free administration (32–34). Vaccines based in oral administration of live *Lactobacillus plantarum* expressing modified OspA_BPBPk_ and intranasal delivery of PIV5-OspA_BPBPk_, respectively, provided 100% protection from tick-transmitted *B. burgdorferi* (11, 31). Yet, the efficacy of parenterally delivered recombinant OspA_BPBPk_, although expected, was not tested previously.

To complete efficacy analysis of our modified OspA vaccine candidate, we asked whether SC prime-boost delivery of OspA_BPBPk_ as well as homologous (IN/IN) vs heterologous (IN/SC) administration routes using intranasal PIV5-OspA_BPBPk_ would improve the magnitude of the antibody response or consistency of this transmission-blocking vaccine. The goal of this study was to test the efficacy of the OspA_BPBPk_ vaccine (11) using different delivery vectors (live PIV5 vs alum adjuvant) and administration routes (intranasal vs subcutaneous) in a tick-challenge murine model of Lyme borreliosis.

## Materials and Methods

### Cells

BHK21 cells (ATCC, Manassas, VA, USA) were cultured in Dulbecco’s Modified Eagle Medium (DMEM; Corning, NY, USA) supplemented with 10% tryptose phosphate broth (TPB; Invitrogen, Thermo Fisher Scientific, Waltham, MA, USA), 5% fetal bovine serum (FBS; Fisherbrand, Pittsburgh, PA, USA), and 1% penicillin-streptomycin (100 IU/mL penicillin and 100 µg/mL streptomycin; Mediatech Inc., Manassas, VA, USA). MDBK cells (ATCC, Manassas, VA, USA) were maintained in DMEM containing 5% FBS and 1% penicillin-streptomycin. All cell lines were incubated at 37°C in a humidified incubator with 5% CO_2_.

### Viruses

The PIV5 plasmid encoding the full-length genome was engineered to express a chimeric *Borrelia burgdorferi/Borrelia afzelii ospA* gene (OspA_BPBPk_) as previously described (11, 31). To rescue the recombinant virus, BHK21 cells at approximately 90% confluence in 6-cm dishes were co-transfected using jetPRIME (Polyplus, Illkirch, France) with the PIV5-OspA_BPBPk_ plasmid along with four helper plasmids: pPIV5-NP, pPIV5-P, and pPIV5-L (encoding the NP, P, and L proteins, respectively), and pT7- polymerase (encoding T7 RNA polymerase). Cells were observed daily for cytopathic effect (CPE) or syncytia, indicating successful virus recovery. The full-length genome of the plaque-purified single clone of PIV5-OspA_BPBPk_ virus was sequenced as previously described (24). Recovered viruses were grown in MDBK cells for 5 to 7 days in DMEM supplemented with 5% fetal bovine serum (FBS) and 1% penicillin-streptomycin (P/S). After incubation, the culture supernatants were collected and centrifuged at 500 × g for 10 min using a Sorvall tabletop centrifuge to remove cellular debris. The clarified virus-containing supernatants were then supplemented with 10% sucrose-phosphate-glutamate buffer (SPG), snap-frozen in liquid nitrogen, and stored at −80°C until use.

### Recombinant OspA_BPBPk_ production

Recombinant protein was expressed in house in *Escherichia coli* BL21 cells grown in Terrific Broth (TB, Fisher BioReagents, Pittsburgh, PA, USA) and purified by Q Sepharose (Cytiva, Marlborough, MA, USA) anion-exchange chromatography. Elution fractions were assessed by Coomassie-stained SDS-PAGE, and fractions showing the strongest target-protein band and fewest visible contaminating bands were pooled. Protein identity was confirmed by Western blotting using anti-OspA monoclonal antibody 184.1. The pooled protein was dialyzed against phosphate-buffered saline (PBS) using a Thermo Fisher Scientific dialysis cassette (Slide-A-Lyzer G3, 10-kDa MWCO; Thermo Fisher Scientific, Waltham, MA, USA), with several changes of 1 L PBS.

### Animals

Thirty-six 8-week-old female C3H-HeN mice were purchased from Inotiv (West Lafayette, IN, USA) and maintained in specific pathogen-free environment at the Laboratory Animal Care Unit of the University of Georgia (UGA) School of Veterinary Medicine with unrestricted access to food and water. Immunized mice were then transferred to the University of Tennessee Health Science Center for tick challenge and assessment of vaccine efficacy. All experiments were performed in compliance with Institutional Animal Care and Use Committee (IACUC) protocols: UGA Protocol no. A2023 01-021-A14 and UTHSC no. 22-0400.

### Experimental design: mouse immunization, challenge and sample collection

Four groups of nine mice (four and five per group in two independent experiments) were anesthetized with isoflurane before receiving the vaccine preparation using a prime– boost schedule on days 0 and 28 (Fig. 1). Briefly, for heterologous vaccinations, mice received a prime intranasal (IN) immunization with 10⁶ PFU PIV5-OspA_BPBPk_ virus in 50 µL administered dropwise to the external nares, followed by a subcutaneous (SC) booster with 20 µg of recombinant OspA_BPBPk_ in 50 µL endotoxin-free PBS (MilliporeSigma, Burlington, MA, USA) combined with 50 µL aluminum hydroxide gel (Alhydrogel; InvivoGen, San Diego, CA, USA), for a final volume of 100 µL (IN/SC). The formulation was prepared approximately 3 h before administration and gently mixed by pipetting. Another set of mice received a prime intranasal immunization with 10⁶ PFU of PIV5 vector control followed by a subcutaneous inoculation of aluminum hydroxide without recombinant protein, formulated as above (IN/SC control). For homologous vaccinations, mice received either intranasal prime and boost with 10⁶ PFU PIV5- OspA_BPBPk_ in 50 µL on days 0 and 28 (IN/IN), or subcutaneous prime and boost, each containing 20 µg recombinant OspA_BPBPk_ in 50 µL endotoxin-free PBS combined with 50 µL aluminum hydroxide gel (SC/SC). Blood was collected on days 27, 63, and 90 by submandibular venipuncture and was allowed to clot for 30 min at room temperature, centrifuged at 2,000 × g for 10 min at 4°C for collection of serum that was stored at −80°C. Serum was used for analysis of anti-OspA_BPBPk_ antibody responses and D90 samples were used for analysis of neutralization of *B. burgdorferi* in BSK-H (Sigma- Aldrich) culture. Mice were challenged on day 90 with nymphal ticks infected with *B. burgdorferi* and all ticks recovered after naturally detaching from the mice were evaluated for presence of *B. burgdorferi flaB* by quantitative polymerase chain reaction (qPCR). Mice were euthanized three weeks after the last day of tick challenge (D120) for collection of samples for downstream analyses (Fig. 1).

**Figure 1.**
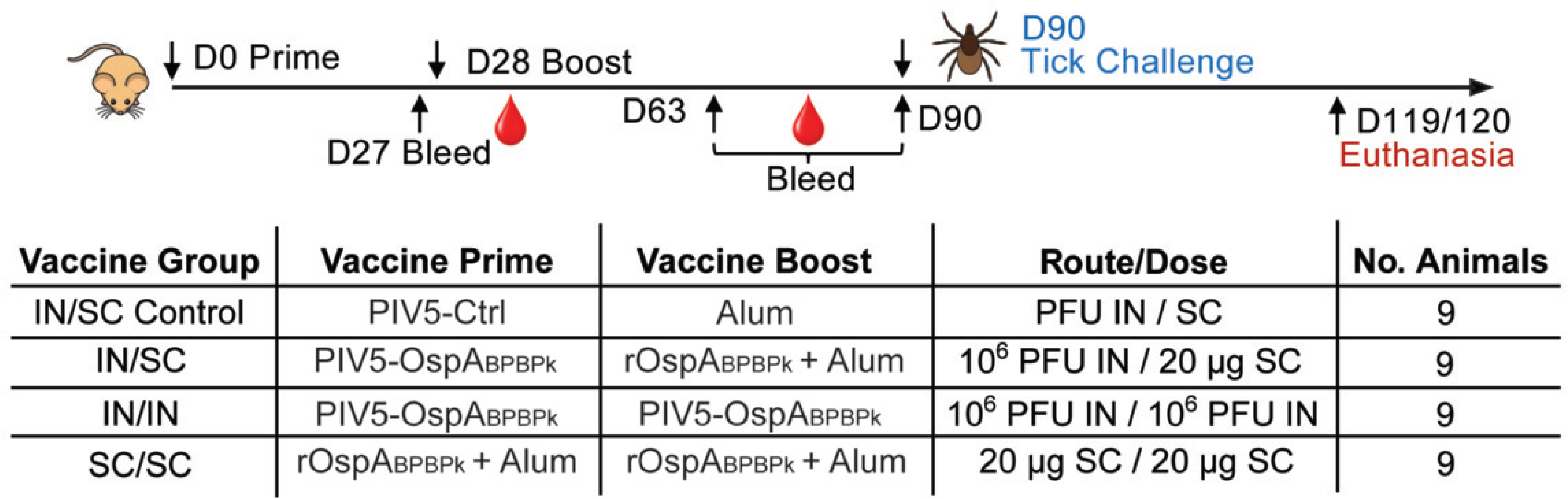
Immunization, sampling, and tick-challenge schedule. Four groups of nine mice received heterologous (IN/SC) and homologous (IN/IN and SC/SC) inoculations on days 0 and 28 in two independent experiments. In the homologous vaccination groups, mice were intranasally (IN) immunized twice (prime/boost) with 10⁶ PFU PIV5- OspA_BPBPk_ (IN/IN) or subcutaneously (SC) with rOspA_BPBPk_ formulated with alum (SC/SC). In the heterologous vaccination groups, mice were intranasally/subcutaneously (IN/SC) immunized with PIV5-OspA_BPBPk_ (10⁶ PFU) / rOspA_BPBPk_ + alum and the negative control received the PIV5 vector + alum (IN/SC control). Collection of blood was done on days 27, 63 and 90. Three weeks after the last day of tick challenge (D120), mice were euthanized for collection of blood, heart, joint, and bladder tissues. Legend: Tick Chlg, tick challenge with ticks carrying 19 strains of *B. burgdorferi*; A_BPBPk_ is modified OspA as described (11, 31).

### Tick challenge

*Ixodes scapularis* nymphal colonies infected with 19 *ospC* types of *B. burgdorferi* were used for challenge. Analysis of strain variability was previously described (11). Briefly, DNA was extracted from five crushed ticks using the NucleoSpin Tissue Kit (Macherey-Nagel, Düren, Germany) and the *ospC* gene was amplified by conventional PCR before being processed for sequencing using the Ion Torrent sequencing instrument (ThermoFisher) at the UTHSC Molecular Resource Center. The infection prevalence in each tick colony was confirmed to be greater than 80% by qPCR targeting the *flaB* gene. On day 90 post-prime immunization, each mouse was individually challenged with 8–10 infected flat nymphal ticks. The fur at the dorsal neck immediately behind the ears was parted, and ticks were placed directly against the skin using fine forceps. Mice were not anesthetized during tick placement and were held individually for < 30 min in a ventilated 50-mL conical-tube restrainer to allow ticks to attach. Each mouse was housed individually in a cage fitted with a grill 2” above a 1” water reservoir and placed within a FIC-2 isolator. Beginning on day 3 after tick placement, the water and cage were inspected daily. Monitoring continued through day 7 to ensure that all ticks placed had engorged and detached, after which mice were returned to their original cages. Detached engorged ticks (> 4 per mouse) were collected from the cage, enumerated, labeled, and stored at –20°C for downstream analysis of *B. burgdorferi* load by *flaB* qPCR. Three weeks after the last day of tick challenge (D120), mice were euthanized by inhalation of 3-5% isoflurane (Sigma- Aldrich, St. Louis, MO, USA) followed by exsanguination via cardiac puncture and thoracotomy; blood, heart, ear, knee joint, and bladder tissues were collected. Bladder and heart tissues were cultured in BSK-H medium (Sigma-Aldrich, St. Louis, MO, USA) to assess spirochete growth and motility by dark-field microscopy, while parallel samples and other tissues were preserved in RNAlater (Invitrogen, Thermo Fisher Scientific, Waltham, MA, USA) for *flaB* qPCR. Serum samples were analyzed for anti-*B. burgdorferi* antibody responses by PepVF ELISA as described below.

### Measurement of antigen-specific antibody

OspA_BPBPk_ and *B. burgdorferi* PepVF- specific antibodies were measured using the enzyme-linked immunosorbent assay (ELISA). 96-well flat-bottom ELISA microtiter plates (Nunc MaxiSorp; Thermo Fisher Scientific, Waltham, MA, USA) were coated overnight at 4°C with 100 µL PBS containing 1 µg/ml OspA_BPBPk_ or 10 µg/ml PepVF (GenScript, Piscataway, NJ, USA). After overnight incubation, the ELISA plate was washed twice with a washing buffer containing 0.01% Tween-20. The plate was blocked by the addition of a blocking buffer containing 1% bovine serum albumin (BSA; Thermo Fisher Scientific, Waltham, MA, USA) followed by incubation for 1 hr at 37°C. After washing, plates coated with PepVF were incubated with serum diluted 1:100. For measurement of anti-OspABPBPk endpoint titers, sera were serially diluted from 1:100 to 1:10⁸. The endpoint titer was defined as the highest serum dilution producing an OD_450_ value above the background cutoff. Plates were then incubated for 30 min with HRP-conjugated goat anti-mouse IgG (H+L; Jackson ImmunoResearch) diluted 1:10,000, followed by standard color development using 3,3′,5,5′-tetramethylbenzidine (TMB) substrate (Rockland Immunochemicals, Limerick, PA, USA). Absorbance was measured at 450 nm using a SpectraMax iD3 plate reader (Molecular Devices, San Jose, CA, USA). PepVF is a synthetic 33-amino-acid *B. burgdorferi* peptide containing sequences derived from the VlsE IR6 region (35) and an internal FlaB fragment and has previously been validated as a sensitive serologic marker of *B. burgdorferi* infection (36). Anti-PepVF responses were therefore measured as evidence of host exposure to *B. burgdorferi* following tick challenge. Seropositivity for anti-PepVF and anti-OspA was defined as an OD_450_ value greater than the mean plus three standard deviations of the negative-control sera.

### *B. burgdorferi* neutralization assay

Neutralization of *B. burgdorferi* was performed as described (11). Briefly, frozen stocks of the same *B. burgdorferi* multi-strain culture used to produce infected ticks (passage 1) were grown to mid-log phase at a density of approximately 1 × 10⁷ motile spirochetes/mL, as determined using a Petroff–Hausser chamber. Mouse sera collected on day 90 from vaccinated and control mice were heat- inactivated at 56°C for 30 min. Eight microliters of *B. burgdorferi* culture was mixed with 4 µL of heat-inactivated serum and 4 µL of reconstituted guinea pig complement used undiluted (MP Biomedicals, Santa Ana, CA, USA) in a 0.2-mL sterile PCR microtube (VWR, Radnor, PA, USA). The positive control group consisted of 8 µL BSK-H medium (Sigma-Aldrich, St. Louis, MO, USA) mixed with 8 µL of *B. burgdorferi* culture. Samples were incubated at 34°C for 6 days. Motile spirochetes were counted in five microscopic fields on days 0, 4, and 6 using a Petroff–Hausser chamber under dark-field microscopy (Axio A1, Zeiss USA, Hawthorne, NY, USA), and the five field counts were averaged for each sample.

### Culture of *B. burgdorferi* from mouse tissues

Heart and bladder tissues collected at necropsy were selected for assessment of disseminated *B. burgdorferi* infection. Approximately half of each heart was placed intact into 5 mL of BSK-H medium, with the remaining half reserved for qPCR, while the whole bladder was placed intact into 5 mL of BSK-H medium. The medium was supplemented with 6% rabbit serum (Sigma-Aldrich, St. Louis, MO, USA) and 1% of a 100× antibiotic mixture containing fosfomycin (Thermo Scientific Chemicals, Waltham, MA, USA), rifampicin (TCI America, Portland, OR, USA), and amphotericin B (MP Biomedicals, Santa Ana, CA, USA), giving final concentrations of 20 µg/mL, 50 µg/mL, and 2.5 µg/mL, respectively. Cultures were incubated at 33°C in closed tubes and examined by a blinded reader under dark-field microscopy on days 14 and 21. Five microscopic fields were examined for each culture, and a culture was classified as positive when motile spirochetes were observed. On day 21, culture supernatant was collected for *flaB* qPCR to support the dark-field microscopy findings and to detect low-level bacterial growth not readily observed microscopically. Detection of *flaB* DNA in culture supernatant alone is not considered evidence of bacterial viability.

### qPCR to quantify *B. burgdorferi* in engorged ticks, mouse tissues and cultures from mouse tissues

Genomic DNA was purified from tick samples and mouse tissues (heart, joint, and ear) using the NucleoSpin Tissue Kit (Macherey-Nagel, Düren, Germany) according to the manufacturer’s instructions. Tick samples were mechanically disrupted in a bead beater (Mini-BeadBeater-16; BioSpec Products, Bartlesville, OK, USA) with 1-mm zirconia beads (BioSpec Products, Bartlesville, OK, USA) before extraction. The purified DNA was eluted in 100 µL of the kit elution buffer and stored at −80 °C for further analysis. For analysis of heart- and bladder-tissue cultures, 2 µL of culture was added directly to the qPCR reaction without prior DNA extraction. Quantification *of B. burgdorferi* was performed using quantitative polymerase chain reaction (qPCR) on a QuantStudio 3 system (Applied Biosystems, Thermo Fisher Scientific, Waltham, MA, USA), targeting the conserved *flaB* gene (11). The primers used were as follows: forward 5′-GCAGCTAATGTTGCAAATCTTTTC-3′, reverse 5′- GCAGGTGCTGGCTGTTGA-3′, and probe 5′-[6-FAM]-

AAACTGCTCAGGCTGCACCGG-[TAMRA-Q]-3′. For generating the standard curve, DNA was extracted from a cultured stock of *B. burgdorferi* with an initial concentration of 10⁶ organisms, as determined by enumeration under dark-field microscopy, and serially diluted from 10⁵ to 1. qPCR reactions were carried out in a final volume of 20 µL using TaqMan™ Fast Advanced Master Mix (Applied Biosystems, Thermo Fisher Scientific, Waltham, MA, USA) containing 900 nM of each primer, 250 nM of the probe, and 2 µL of DNA template. Duplicate reactions were performed. The amplification protocol included an initial step of 10 min at 95°C, followed by 40 cycles of amplification (15 s at 95°C and 1 min at 60°C). For tissue samples, DNA was extracted from 25 mg of tissue, and flaB copy numbers were calculated and reported per milligram of tissue. For tick samples, *flaB* copy numbers were reported per individual whole tick.

### Statistical analysis and Reproducibility

All statistical analyses were performed in GraphPad Prism version 10.6.1 (GraphPad Software, Boston, MA, USA) and Python version 3.14.6 with SciPy version 1.18.0. Sample sizes (n = 9 mice per group) were based on prior power calculations and are standard for controlled tick-challenge studies. Other datasets were analyzed using tests matched to their distributional properties, as specified in the figure legends. Anti-OspA_BPBPk_ ELISA endpoint titers (Fig. 2) were analyzed using the Kruskal–Wallis test with Dunn’s multiple-comparisons post hoc test. Tick *flaB* qPCR copy-number data (Fig. 3A–B) were analyzed using a mixed-effects model with immunization group as a fixed effect and experiment as a random effect.

**Figure 2.**
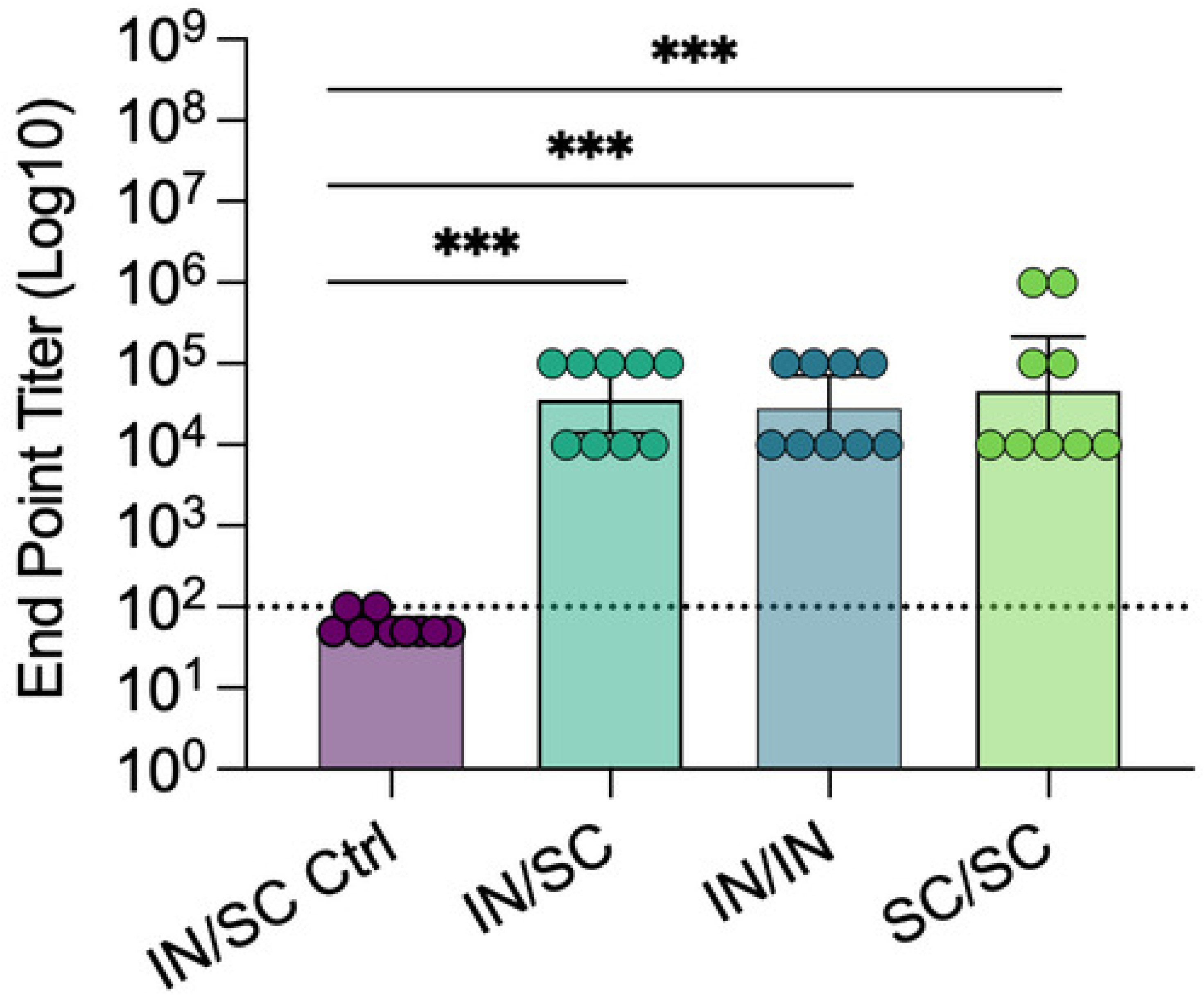
Homologous and heterologous OspA_BPBPk_ vaccination elicits high IgG endpoint titers. Anti-rOspA_BPBPk_ total IgG endpoint titers were measured by enzyme- linked immunosorbent assay (ELISA) using serum serially diluted from 1:100 to 1:10⁸. The endpoint titer was defined as the highest serum dilution with an OD₄₅₀ value above the background cutoff. Each point represents one mouse, and bars show the geometric mean with 95% confidence interval. Statistical significance was determined using the Kruskal–Wallis test with Dunn’s multiple-comparisons test. \*\*\**p* < 0.001. IN/IN, intranasal prime and boost; SC/SC, subcutaneous prime and boost; IN/SC, intranasal prime and subcutaneous boost; IN/SC control, intranasal control-vector prime and subcutaneous adjuvant-control boost.

**Figure 3.**
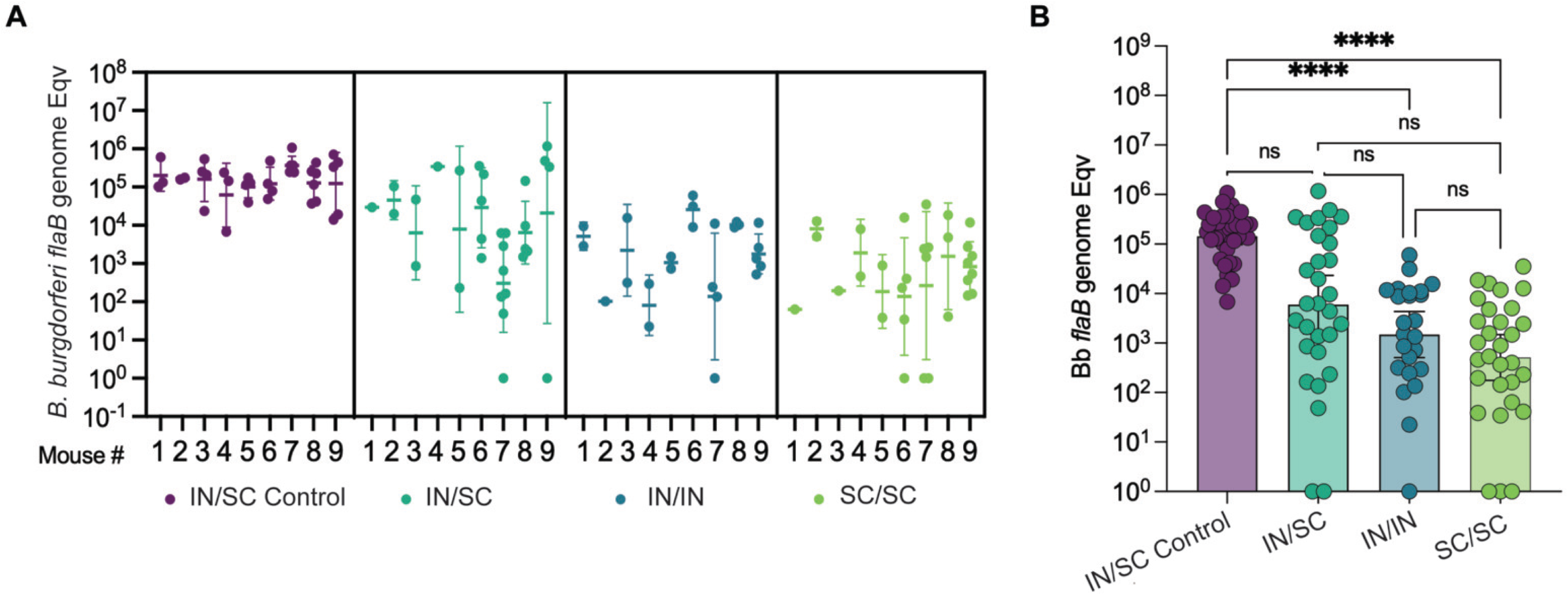
Homologous and heterologous OspA_BPBPk_ vaccination reduces *B. burgdorferi* burden in engorged ticks after feeding. *B. burgdorferi* flaB genome equivalents were quantified by qPCR in individual engorged nymphal ticks recovered after feeding. (A) Tick burdens are shown according to the individual mouse on which each tick fed; each point represents one tick, and horizontal bars indicate the geometric mean. (B) Tick burdens are shown by vaccination group; each point represents one tick, and bars indicate the geometric mean. Statistical significance was determined using a mixed-effects model with immunization group as a fixed effect and experiment as a random effect, followed by Dunnett’s multiple-comparisons test. \*\*\*\**p* < 0.0001; ns, not significant; Eqv, genome equivalents. IN/IN, intranasal prime and boost; SC/SC, subcutaneous prime and boost; IN/SC, intranasal prime and subcutaneous boost; IN/SC control, intranasal control-vector prime and subcutaneous adjuvant-control boost.

Neutralization assay data (Fig. 4A–B) were analyzed within each experimental group using the Friedman test for matched observations across days 0, 4, and 6, followed by Dunn’s multiple-comparisons test comparing days 4 and 6 with day 0. Anti-PepVF ELISA data (Fig. 5A), *flaB* in tissues qPCR (Fig. 5B) and culture-enriched *flaB* qPCR data (Fig. 5C) were analyzed by 2-way ANOVA. Biological replicates were individual mice, each contributing multiple ticks and tissues, and findings were reproduced across two independent experiments.

**Figure 4.**
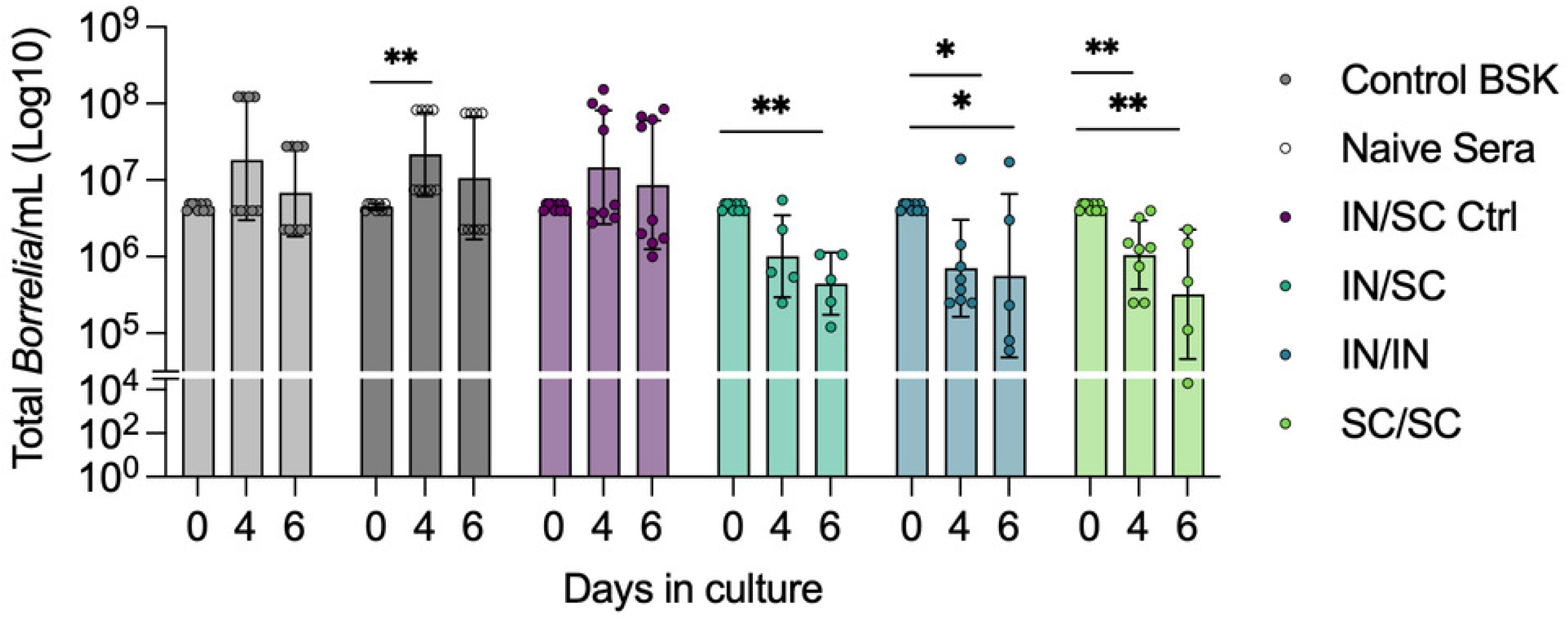
Sera from homologous and heterologous vaccination regimens exhibit anti-*B. burgdorferi* neutralizing activity. Sera collected from experimental mice (IN/IN, SC/SC, IN/SC and IN/SC control) before tick challenge (D90) were incubated with a multi-strain culture of *B. burgdorferi* for up to 6 days. Additional control groups included BSK-H medium and naïve serum. Samples were collected on days 0, 4, and 6 and motile spirochetes were quantified under a dark-field microscope. Scatter plots (A– B) represent numbers of motile *B. burgdorferi* (geometric mean with geometric standard deviation [SD]). Each dot represents data from a single mouse. Statistical significance was determined within each group using the Friedman test followed by Dunn’s multiple- comparisons test comparing days 4 and 6 with day 0. Only significant comparisons are shown;\**p* < 0.05 and \*\**p* < 0.01.

**Figure 5.**
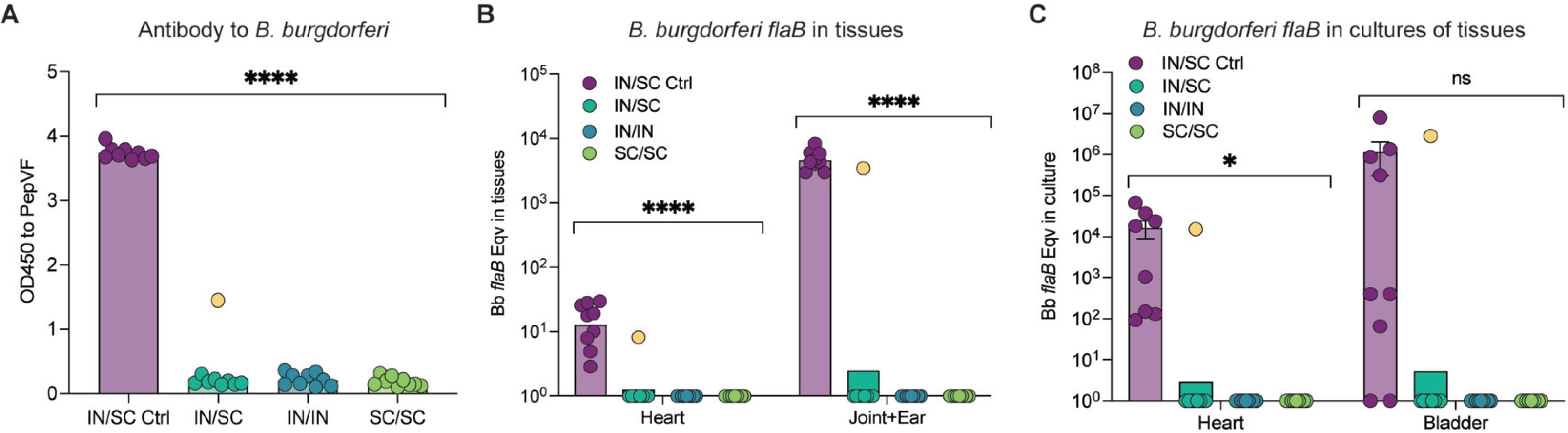
Homologous and heterologous vaccination with prevents dissemination of *B. burgdorferi* to target tissues three months post-prime vaccination. Scatter plots representing geometric mean of *B. burgdorferi* anti-PepVF antibody measured by ELISA (A), *B. burgdorferi flaB* burden in heart, joint, and ear tissues quantified by qPCR and reported as copies per milligram of tissue (B) and *B. burgdorferi flaB* in 4-week cultures of heart and bladder tissues quantified by qPCR (C). Each dot represents data from a single mouse, yellow represents the same mouse across panels; n=9 per group; Statistics by 2-way ANOVA, **** *p*<0.0001; * *p*=0.0182; ns *p*>0.05.

## Results

### Antibody responses following vaccination

All OspA_BPBPk_-containing vaccination regimens (IN/IN, SC/SC, IN/SC) elicited comparable antigen-specific IgG endpoint titers among groups, with geometric mean titers (GMTs) of approximately 4–6 log₁₀, which were significantly higher than the IN/SC vector controls of ∼2 log₁₀ (*p* < 0.001). These results demonstrate that both homologous and heterologous vaccination strategies induced strong systemic IgG responses against OspA_BPBPk_ (Fig. 2).

### Impact of OspA vaccination on *B. burgdorferi* in feeding ticks

Engorged ticks collected after natural detachment from homologous IN/IN and SC/SC mice carried significantly fewer *B. burgdorferi* organisms than those from control animals (IN/SC control) (*p* < 0.0001 for both groups), whereas the heterologous IN/SC group showed a reduction that was not significant relative to control (*p* = 0.2456) (Fig. 3). Quantitative *flaB* qPCR showed a geometric mean burden of 1.78 × 10⁵ copies in the IN/SC control group, compared with 3.77 × 10³ copies in the heterologous IN/SC group and 2.56 × 10³ and 1.76 × 10³ copies in the homologous IN/IN and SC/SC groups, respectively. These values correspond to a 1.7 log₁₀ reduction in the heterologous regimen and 1.8–2.0 log₁₀ reductions in the homologous regimens. Differences were not significant between vaccinated groups. Mixed-effects model analysis and multiple-comparisons testing are shown in Fig. 3B.

### Neutralization of *B. burgdorferi* motility

To determine whether vaccination elicited functional anti-*B. burgdorferi* activity, D90 sera collected before tick challenge were evaluated in an *in vitro* neutralization assay by counting motile *B. burgdorferi* spirochetes under dark-field microscopy at days 0, 4, and 6 (Fig. 4). Sera from both homologous vaccination regimens showed significant reductions in motile spirochetes relative to D0 at both time points tested (IN/IN at D4 with *p* = 0.0454 and D6 with *p* = 0.0386, SC/SC at D4 with *p* = 0.0037 and D6 with *p* = 0.0015). In the heterologous IN/SC group, the reduction at D4 was not significant (*p* = 0.1122), but a significant reduction was observed at D6 (*p* = 0.0048). In contrast, numbers of motile spirochetes in control groups (BSK-H medium, naïve serum, and IN/SC control serum) remained similar or increased (D4 naïve serum). These findings indicate that both homologous and heterologous vaccination regimens elicited functional anti-OspA_BPBPk_ antibodies that neutralized *B. burgdorferi* motility in culture, with the homologous regimens showing earlier detectable neutralizing activity in this assay.

### Prevention of *B. burgdorferi* dissemination to host tissues and overall vaccine efficacy

To assess *B. burgdorferi* dissemination and systemic infection following tick challenge, we quantified anti-PepVF antibody responses in serum from vaccinated mice, *B. burgdorferi flaB* DNA in tissues and *flaB* DNA in cultures from tissues (Fig. 5). Dark-field microscopy (DFM) observation of live *B. burgdorferi* in cultures of heart and bladder tissues collected after euthanasia was used as the primary measure of overall vaccine efficacy (Table 1). In contrast to the control, no evidence of *B. burgdorferi* dissemination or systemic infection with live spirochetes was observed in any mice in the IN/IN and SC/SC groups, as shown by absence of antibody to *B. burgdorferi* PepVF (Fig. 5A), *flaB* DNA in heart, joint or ear tissues or *flaB* DNA in cultures of heart and bladder tissues (Fig. 5B–C). In the heterologous IN/SC group, one out of nine mice (1/9, 11%) had anti- *B. burgdorferi* PepVF antibody in serum (Fig. 5A), and *flaB* DNA was present in heart, joint or ear as well as in supernatant from heart and bladder tissue cultures (Fig. 5B–C). Regarding observation of vigorously motile *B. burgdorferi* in cultures of tissues by DFM (Table 1), no motile spirochetes were recovered from cultures of heart and bladder from IN/IN, SC/SC and IN/SC vaccinated groups, which was in contrast to the IN/SC control that showed vigorously motile spirochetes in one or both tissues. Absence of live spirochetes in tissues from the IN/SC vaccinated mouse indicates absence of active systemic infection. These results confirm that OspA_BPBPk_ prime-boost vaccination effectively prevents active systemic *B. burgdorferi* infection following tick challenge independently of vaccine delivery vehicle (PIV5 or alum adjuvant) and administration route (intranasal or subcutaneous) in mice.

**Table 1.**
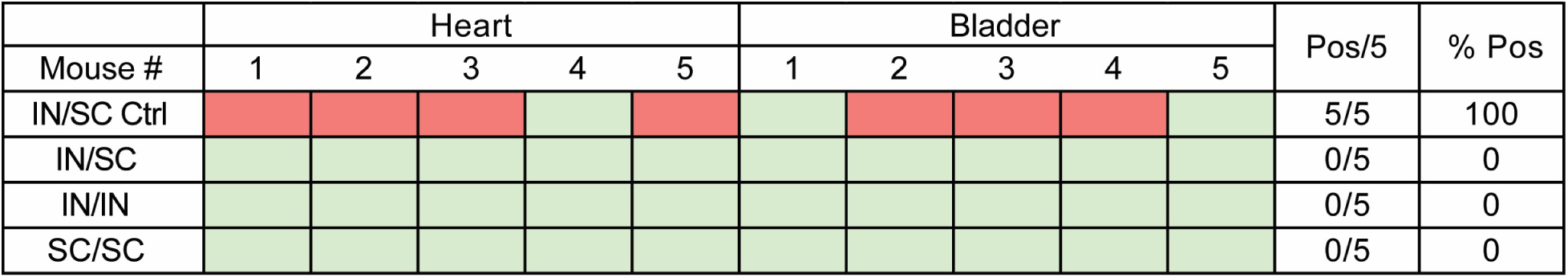
Vaccine efficacy. Presence of motile spirochetes was determined by dark-field microscopy after 21 days of culture of heart and bladder tissues in BSK-H medium. A mouse was classified as culture positive if either heart or bladder culture yielded motile organisms. This group of samples (n=5 per group) included tissues from the IN/SC mouse that produced positive PepVF and *flaB* qPCR results. Legend: IN/SC control, intranasal control-vector prime and subcutaneous adjuvant-control boost; IN/SC, intranasal prime and subcutaneous boost; IN/IN, intranasal prime and boost; SC/SC, subcutaneous prime and boost.

## Discussion

Both delivery vehicles and all routes of administration elicited strong systemic anti- OspA_BPBPk_ IgG responses in mice, consistent with previously reported protective anti-OspA titers. Murine tissue *B. burgdorferi* burdens were mostly absent across all vaccinated groups, with homologous regimens providing the most consistent outcomes across serologic, molecular, and culture-based measures. A single heterologous vaccinated mouse had *flaB* DNA in tissues after tick challenge, although no motile organisms were observed in cultures of the same tissues, suggesting dissemination of *B. burgdorferi* without evidence of recoverable viable infection. Overall, repeated delivery via the same route produced more uniform antibody responses in mice, although the underlying mechanism was not fully examined here.

Heterologous prime-boost strategies, in which different vaccine modalities or routes are combined, are often used to broaden immune responses, enhance durability, or augment mucosal immunity by engaging complementary immunologic mechanisms, particularly for respiratory and enteric pathogens (37, 38). For OspA-based Lyme disease vaccines, however, protection is determined primarily by the concentration of circulating anti-OspA antibodies reaching the tick midgut during engorgement (16). Specifically, antibodies acquired from vaccinated hosts act within the engorged tick midgut to neutralize spirochetes before migration of *B. burgdorferi* to the salivary glands occurs (13, 14, 39). Thus, the rationale for testing the heterologous IN/SC regimen was not to enhance mucosal protection, but to determine whether combining an intranasal PIV5-vectored prime with a recombinant OspA_BPBPk_ protein delivered in alum could improve the magnitude or consistency of the antibody response mediating transmission blocking while maintaining needle-free delivery of one of the two doses. Therefore, unlike mucosal-entry pathogens where compartmentalization can be critical, the present data support the interpretation that protection in this model is associated with achieving high systemic anti-OspA IgG responses.

Both homologous and heterologous regimens generated high systemic IgG titers and conferred protection after tick challenge, but the regimens were not identical by every measure. The homologous regimens (IN/IN and SC/SC) were associated with the lowest tick *flaB* burdens. In addition, sera from the homologous regimens showed earlier detectable neutralizing activity of *B. burgdorferi* motility (day 4), whereas the heterologous IN/SC regimen showed a later effect (day 6) (Fig. 4). This result correlated with *B. burgdorferi* dissemination as we detected *B. burgdorferi flaB* DNA in tissues from one mouse in the IN/SC group (Fig. 5) in contrast to homologous IN/IN and SC/SC groups. Nevertheless, the ultimate protection outcome in this study (absence of culturable, motile *B. burgdorferi* in tissues) did not differ among the vaccinated groups. We also show that subcutaneously delivered recombinant OspA_BPBPk_ adjuvanted with aluminum hydroxide protected against tick-transmitted *B. burgdorferi*. These results extend our prior work on administration of Lyme disease vaccines. We previously demonstrated protective efficacy of the modified OspA_BPBPk_ vaccine candidate using an oral *Lactobacillus plantarum* delivery system (31) and showed that intranasal PIV5-OspA_B31_ and PIV5-OspA_BPBPk_ vaccination provided durable (>1 year) protection against tick-transmitted *B. burgdorferi* (11). The comparable efficacy of heterologous and homologous regimens in prevention of culturable infection in mice is consistent with the view that systemic antibody levels, rather than route diversification, are the main predictor of protection in OspA-based transmission-blocking vaccines (13, 14, 17). Comparable correlations between antibody titer and protection have been reported for recombinant OspA vaccines (3, 4) and for more recent multivalent formulations such as VLA15, in which passive-transfer experiments established antibody thresholds of ∼130 U/mL sufficient to prevent transmission (18). Together, these findings suggest that maintenance of high and uniform systemic anti-OspA IgG responses is more important than diversifying immunization routes.

Durability of protection beyond 90 days post-prime remains to be determined. Future work should extend regimens to longer intervals, evaluate dose de-escalation regimens to determine whether heterologous delivery offers an advantage when antibody responses are lower, evaluate multivalent OspA formulations to capture genospecies diversity (1, 40) and characterize antibody durability and functional neutralization as correlates of protection. Another limitation of the current study was that we did not assess sex as a biological variable. Female mice were used to maintain consistency with prior experiments with the established tick-challenge model (41).

In conclusion, both homologous and heterologous prime–boost regimens using recombinant OspA_BPBPk_ and PIV5-OspA_BPBPk_ vaccines elicited strong systemic antibody responses and protected mice against tick-transmitted multi-strain *B. burgdorferi* comprising 19 OspC types. While heterologous delivery performed comparably, homologous intranasal and subcutaneous regimens provided the most consistent outcomes as assessed by culture of *B. burgdorferi* from tissues after tick challenge. These results support the view that high and sustained systemic IgG titers to OspA are closely associated with effective transmission blocking in this model. The PIV5 vector platform is undergoing further testing for future Lyme disease vaccine development and may help inform simplified, dose-sparing immunization strategies for human application.

## Author Contributions

MCG, BH and MGS were responsible for the concept and design of the experiments. MCG, OA, MJS, KBH conducted the assays. MCG, GJ, SK and MGS analyzed the data. BH, HJ and MGS secured funding, administered the collaborative project, and supervised personnel. MCG, GJ and MGS wrote the manuscript.

## Funding

This work was supported by the National Institute of Allergy and Infectious Diseases (NIAID), United States National Institutes of Health (NIH), grant numbers R01 AI139267 (MGS), R44 AI167605 (MGS) and funding from CyanVac, LLC. BH has been supported by Fred C. Davison Distinguished University Chair in Veterinary Medicine. The content is solely the responsibility of the authors and does not necessarily represent the official views of the NIH.

## Acknowledgements

We are grateful to our colleagues and collaborators for helpful discussions and comments on the manuscript. We also thank the Ae Kyung Yi laboratory for access to imaging equipment used for Western blot documentation (Chemi-Doc).

## Competing Interests

MGS is President and CEO of Immuno Technologies, Inc. BH is President and CEO of CyanVac, LLC. MCG, HJ, BH and MGS hold a patent that covers use of PIV5-OspA-based vaccines. BH, MCG and HJ are employees of CyanVac, LLC and hold equity in its affiliate. The other authors have no competing interests.

## Notes

### Summary of Updates

This version has been revised in response to reviewer comments. The Introduction and Discussion were updated to clarify the mechanism of OspA mediated protection, the role of circulating anti OspA antibodies, and the rationale for homologous and heterologous vaccination. The Methods were expanded to provide additional detail on vaccine preparation, PIV5 production, tick challenge, sample collection, culture, qPCR, ELISA, neutralization, and statistical analysis. The Results and Discussion were revised to more clearly distinguish the outcomes of the different vaccination regimens and to clarify interpretation of the single heterologous vaccinated mouse with evidence of exposure. Figure 1 was revised to improve clarity, figure legends were updated, and the reference list was expanded and corrected.

